# A Pipeline for Solving Edge-Matching Puzzles and Their Implications for Protein Folding

**DOI:** 10.64898/2026.05.23.727379

**Authors:** Shahar Seifer

**Affiliations:** Chemical and Biological Physics, Weizmann Institute of Science, Rehovot 761001, Israel

**Keywords:** Eternity II puzzle, quantum annealing, adiabatic quantum computing, protein folding

## Abstract

Progress in quantum computation offers new opportunities for addressing longstanding combinatorial challenges. One such challenge is the Eternity II edge-matching puzzle, consisting of 256 tiles, which has resisted solution despite extensive community effort. The computational complexity of this NP-complete problem exceeds the capacity of current quantum annealing processors but lies within reach of hybrid quantum-classical solvers. Testing a quadratic unconstrained binary optimization (QUBO) model of a puzzle on a D-Wave hybrid solver demonstrates a complete solution only for puzzle instances up to 64 tiles. Simulated quantum annealing fails on this benchmark, whereas an original classical heuristic, “nucleation with deduction”, succeeds. To approach the full Eternity II puzzle, I developed a MATLAB package that integrates multiple quantum and classical approaches, including neural-network transformers and gradient-based refinement. A multistage computation pipeline is demonstrated successfully on a puzzle comparable in complexity to Eternity II and with a known solution, based on multiple hybrid optimization steps with both “hard” and “soft” constraint formulations, identification of persistent substructures, and a final classical refinement stage. The resulting optimization problem involves ∼100,000 logical variables and requires partial initialization. Intriguingly, solving this puzzle mirrors the “end game” of protein folding, a process that nature completes in mere fractions of a second, seemingly defying expectations set by the Levinthal paradox. The prospect of predicting protein structure by quantum annealing is reviewed in light of these results.

## I. INTRODUCTION

An NP-complete problem is one whose computational complexity grows faster than any polynomial function of its input size. It is termed nondeterministic polynomial (NP) because a proposed solution can be verified in polynomial time. Once a solution method is found for one NP-complete problem, even one arising from a game, it applies to all others through appropriate transformations [1], [2]. Edge-matching puzzles, in particular, offer an instructive example of how complexity grows exponentially when partial solutions cannot be validated [3]. These differ from common jigsaw puzzles, whose pieces match unambiguously and therefore exhibit minimal frustration.

The Eternity II puzzle is one such difficult challenge that I have attempted to solve in this work, despite its careful design by Christopher Monckton to eliminate hidden shortcuts. A $2 million reward offered between 2007 and 2010 remained unclaimed, and a dedicated community of solvers continues to pursue this challenge [4]. Notable academic contributions include a proof that look-ahead strategies are ineffective [5], as well as a PhD dissertation [6]. The highest score achieved in the Eternity II competition was 467 out of 480 matching edges by Louis Verhaard. Improvements in classical computation have since reached a maximum score of 470 out of 480.

The Eternity II puzzle comprises 256 square tiles (also referred to as pieces), each featuring four edge patterns that must be matched with adjacent tiles. With 23 distinct edge types, the puzzle offers a compelling analogy to a typical protein, which is built from up to 22 standard amino acids. The difficulty is evident when considering partial solutions shared by the solver community: these partial solutions rarely overlap and likely do not resemble the final solution. For example, the runner-up of the 2010 competition later generated two additional partial solutions with similar scores [7], and comparison between them shows no more than one or two pieces in common. A full solution is said to exist but has not been revealed.

A binary representation of the problem follows from the 16×16 positions of square pieces, each with four possible orientations. Consequently, each position initially has 1,024 possible assignments (piece plus orientation). Eliminating options for edge pieces and incorporating initial clues results in roughly 100,000 logical variables representing available selections. The daunting task of finding a complete solution is equivalent to navigating a multi-dimensional maze in which each degree of freedom represents a decision point. The benefit of quantum computation is akin to “seeing through the walls” of this maze. In D-Wave quantum annealing hardware [8], the binary options 0 and 1 are represented by a spin state *S*_*i*_, which upon measurement are selected between +1 and −1 (spin up or down). The quantum annealer returns a spin configuration {*S*_*i*_}that minimizes the energy *H* of the Ising spin model.

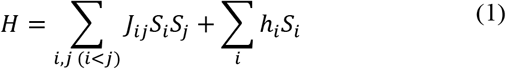

where the coupling interactions *J*_*ij*_ and magnetic fields *h*_*i*_ are programmable inputs. In a quantum annealing computer, the coupling is gradually activated, typically over milliseconds, allowing the system’s final spin configuration to approach a minimum-energy state. This process is often described as quantum tunneling, enabling transitions between local minima rather than becoming trapped in suboptimal configurations. Achieving the global minimum represents an NP-hard challenge, whereas settling into a local minimum is a classical problem commonly addressed by machine-learning algorithms.

However, current technology has serious limitations: (a) Quantum annealing cannot guarantee that a returned solution is the global minimum; (b) a quantum processing unit contains up to ∼5,000 spins, each limited to ∼24 couplers; and (c) The effective resolution of coupling coefficients is 4–5 bits (according to D-Wave support). These constraints imply that an effective NP-hard solver requires a hybrid approach combining quantum and classical computation. The D-Wave hybrid solver partitions the QUBO into subproblems that are either fitted to hardware or attempted classically [9].

Alternative classical approaches for solving QUBO problems are included in software packages such as OpenJij [10], Gurobi, CPLEX, CBC, and components of the D-Wave Ocean library. Strategies such as branch-and-bound or integer linear programming guarantee optimality but scale poorly, typically failing beyond a few thousand binary variables unless the structure is highly sparse. Classical heuristics, including simulated annealing, tabu search, large-neighborhood search, and evolutionary methods, scale to larger instances but provide no guarantees and often struggle in highly constrained landscapes. For the scale of the Eternity II puzzle no classical method competes effectively with the hybrid quantum annealer as a standalone strategy.

It is possible to reduce problem dimensionality slightly by identifying persistent variables. These are QUBO variables that remain stable in value when solved for “relaxed” versions of the problem with polynomial complexity [11]. The method is most effective when persistent variables can be determined in advance. Yet, as shown in the results section, incomplete solutions produced by a hybrid solver also reveal persistent variables that cannot be known beforehand but can nevertheless be identified across repeated trials.

The work presented here considers hybridization between quantum annealing steps based on a QUBO formulation and classical algorithms specifically efficient for solving edge-matching puzzles. Both approaches offer complementary strengths, and an efficient hybrid pipeline succeeds in recovering a complete solution. In the final section, I address a fundamental scientific challenge of determining the structure of a protein from its sequence and argue for similar hybrid strategy.

## II. METHODS

The following techniques were implemented in several Matlab scripts and python modules used to explore solution pipelines for edge-matching puzzles. The code and results are publicly available at https://github.com/Pr4Et/EdgeMatchingPuzzleSolver.

### A. QUANTUM ANNEALING APPROACH

The QUBO problem is formulated in an Ising model representation, where each spin encodes an available option for placing a specific piece in a specific orientation at a given board position. A spin value of *S* =−1 (down) denotes selection of that option. The configuration space assumes that for every possible placement of a piece, there exist potential matches with adjacent pieces.

Consider adjacent board positions A and B, and let k denote any possible symbol that can appear on their shared edge. between the pieces. The sets of allowed (piece, rotation) options for this symbol are denoted by *A*_*k*_ and *B*_*k*_ respectively. In a correct solution, the number of spins down in *A*_*k*_ and *B*_*k*_ must match. Thus, we minimize

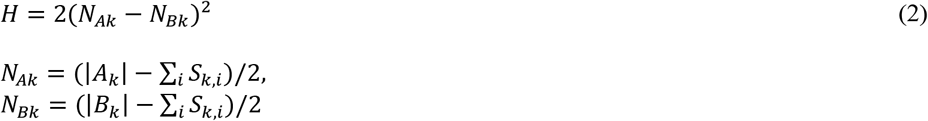

|*A*_*k*_| denotes the number of spins in group *A*_*k*_. The Ising model enforces this condition through the energy expression of Eq.1, with the following linear coefficients for each spin *i* in *A*_*k*_ or *B*_*k*_

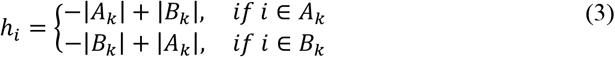

and quadratic couplings for each spin pair *i,j* in the matching groups

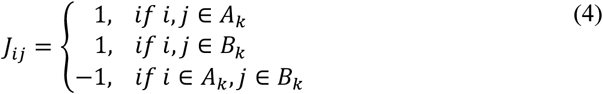

This construction is repeated for each pair of adjacent board positions, producing multiple quadratic terms in *H*. Because each spin participates in up to four independent matching groups, the resulting structure is highly constrained and complex.

A valid solution must additionally ensure that each physical piece is used exactly once. One constraint enforces that the total number of selected options at any location equals 1, meaning

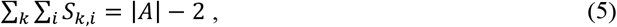

where |A| is the number of spins (≥ 1) associated with that position A. Thus, we add a small penalty term (α<<1) via Eq.1 with coefficients

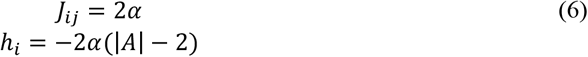

However, due to memory limitations and hybrid-solver efficiency, this constraint is only applied to board locations with fewer than ∼100 options, reducing the total number of coupling terms.

A second constraint ensures that each physical piece appears only once across the entire board. Namely, the same piece cannot be chosen at two different positions nor different orientations simultaneously. A unique solution can be achieved with contribution factor of β=0.25. Thus, the following added terms are written for a group of M spins that represent the same physical piece

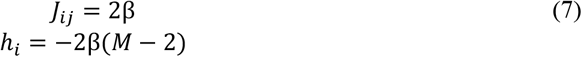

In especially complex instances, enforcing uniqueness strictly can overwhelm the hybrid solver. In such cases, a partial solution with multiple open options can be more useful than a unique but incorrect configuration. To generate such an open partial solution, Eqs.5-7 are replaced by a soft constraint that avoids the quadratic penalty for non-unique solution. Within the limitation of the Ising model, this can be accomplished by the following ancilla method. The M spins for a piece are split into two equal groups. If M is odd a constant spin *S*_*M*+*δ*_ = 1 is added, where δ=1; otherwise δ=0. One ancilla spin *S*_*anc*_ is added per piece. The soft constraint contributes

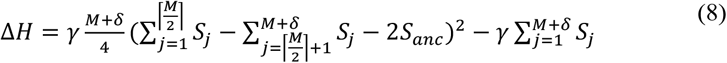

Further details supporting Eq.8 and the derivation of Eqs.3-4 are provided in the Supplementary Material.

### B. CLASSICAL TOOLS: SELECTION WAVES, DEDUCTION, AND NUCLEATION

The propagation of information in the configuration space of the puzzle is described as “selection waves” initiated by selecting pieces that influence the rest of the board. These selections may be integrated in a simplified manner when correlations between non-adjacent positions are ignored. This approach is analogous to the classical Green-function method, which constructs general solutions by integrating over simpler component solutions.

To illustrate the approximation involved it is instructive to consider first the traveling salesman path problem (TSPP). The problem is represented as a graph whose nodes correspond to cities and edges correspond to roads. Each edge is assigned a weight equal to the travel distance *d*_*ij*_ between cities *i* and *j*. The task is to visit all cities exactly once in an order that minimizes total travel distance, finishing at any city (not necessarily the starting point). The Ising model formalism for TSPP is provided in the Supplementary Material. An exhaustive solution searches all possible paths and records their histories, resulting in exponential information growth. Equivalently, one may find the shortest path by propagation information with delays proportional to travel distances. When a simulated message reaches a city, the city’s index is appended and the updated message is transmitted along all outgoing edges. The optimal path is the earliest message containing all node labels. This search cannot be accomplished in polynomial time. However, discarding the combinatorial history allows an approximate solution.

Selection Waves is a reduction of the above to polynomial complexity by retaining only the leading message per time step. If two messages arrive at the same node simultaneously, one is artificially delayed by adding one time unit to its edge delay, ensuring unique arrival times. Particularly for solving edge-matching puzzles, the method follows similar rules by constructing a path along matched pieces. However, here the delay values depend on the accumulated history of each path because the matching score depends on previously visited pieces. Relaxing this dependency restricts the process to a limited set of propagation patterns, producing a partial rather than exhaustive solution. The resulting output is a partial puzzle configuration based on the most favorable propagation path.

Deduction is another form of selection waves that do not retain path history, because it is not needed. A purely deductive process proceeds by eliminating incompatible options based on exposed pieces, which then impose additional constraints across the board. The outcome is an updated configuration space that still contains the correct full solution, provided the initial setup was valid.

The deductive process unfolds iteratively: constraints arriving from any direction refine the allowed pieces at each location, and these refinements propagate outward, eliminating incompatible options elsewhere. Although this elimination does not rely on deep combinatorial reasoning, it substantially reduces problem complexity. If all options at a location are eliminated, the selection waves stall, indicating incorrect initial placements. Thus, a key challenge is providing the solver with a sufficiently accurate initial configuration.

Nucleation is the most direct classical strategy implemented in the software. It operates with full deduction at each step. Nucleation performs an exhaustive search to grow a consistent assembly of matching pieces. Deduction fills in pieces that must follow from earlier selections. If stalling occurs, the process backtracks to earlier positions and explores alternative choices. Attempts to accelerate progress using gradient descent on a pseudo-score function were tested but found ineffective.

### C. PERSISTENCE SEARCH AND NEURAL NETWORK ASSISTANCE

A delicate stage of the solution pipeline is sorting out reliable content from the partly incorrect results obtained from the hybrid quantum solver. The code supports multiple strategies for extracting confident results from several instances. A straightforward approach is identifying persistent placements across many partial solutions. Another approach uses a neural network trained to identify reliable subsets within these partial solutions. Artificial intelligence can sometimes generalize deeply, an ability known as “grokking” [12], suggesting that a model may learn relationships among partial solutions and detect persistent connections that are more likely to assemble into a complete solution.

The implemented AI workflow uses a grid-to-grid transformer architecture. It is trained to analyze connections between pieces (with fixed orientations) and classify each connection as either inside or outside a complete solution. Training used one million randomly generated 16×16 puzzles, from which connection maps were extracted and augmented with randomly generated fake connections of increasing complexity. A directional-attention encoder captured local and pairwise structure, and a transformer-based decoder predicted the likelihood that each connection belonged to a valid solution. In simulation, the model correctly identified 99% of true connections even when up to 1,200 fake connections were added. To address the issue of uncertain orientations, the analysis is repeated over several passes, each time fixing a randomly chosen orientation and ignoring the others. The final predicted set of pairwise connections is then consolidated into a single board configuration.

## III. RESULTS AND DISCUSSION

A straightforward implementation of the D-Wave hybrid solver is first demonstrated on an 8×8 board. Next, a 16×16 board, referred to as the “virtual” puzzle, is examined. This puzzle is comparable in complexity to the Eternity II puzzle and uses the same set of clues, except that its full solution is known in advance. The layout of the virtual puzzle was generated from a partial Eternity II configuration, with mismatched pieces adjusted to produce a consistent final arrangement. Output files for both the simulated puzzles and the original Eternity II puzzle are available in the GitHub project repository.

### A. QUANTUM ANNEALING IN MODERATE SIZE PUZZLES

An 8×8 puzzle was generated by randomly assembling edge-matched pieces. The MATLAB script received the inventory of pieces along with several pre-exposed pieces as input. A spin was defined for every potential placement of a piece in a specific orientation at a specific location. The corresponding Ising model was configured according to Eqs.2–3 and Eqs.4–6, enforcing hard constraints using α = 0.0125 and β = 0.25. Under this formulation, simulated quantum annealing using the Ocean library repeatedly failed to find a solution, determining less than 50% of the pieces. In contrast, the D-Wave hybrid binary-quadratic solver (version 2p) produced a complete solution.

Fig.1a shows the resulting complete solution obtained by the hybrid solver using 1576 spins and 4.7 seconds of computation, beginning from 28 pre-exposed frame pieces. A similar solution produced by classical nucleation required roughly 2 minutes. Fig.1b shows a partial solution provided by the hybrid solver for the same 8×8 puzzle, starting from a configuration of 4 corner pieces. The problem required 4,688 spins and a 14-second run time. A total of 83 out of 84 edges were correctly matched. The faults, marked by orange crosses, are due to repeated use of the same piece. There are 9 persistent pieces—shared non-trivially between the partial and complete solutions—marked with orange stars in the figure. Increasing the contribution factor β slightly, to 0.27, was less successful in terms of matching edges.

**FIGURE 1.**
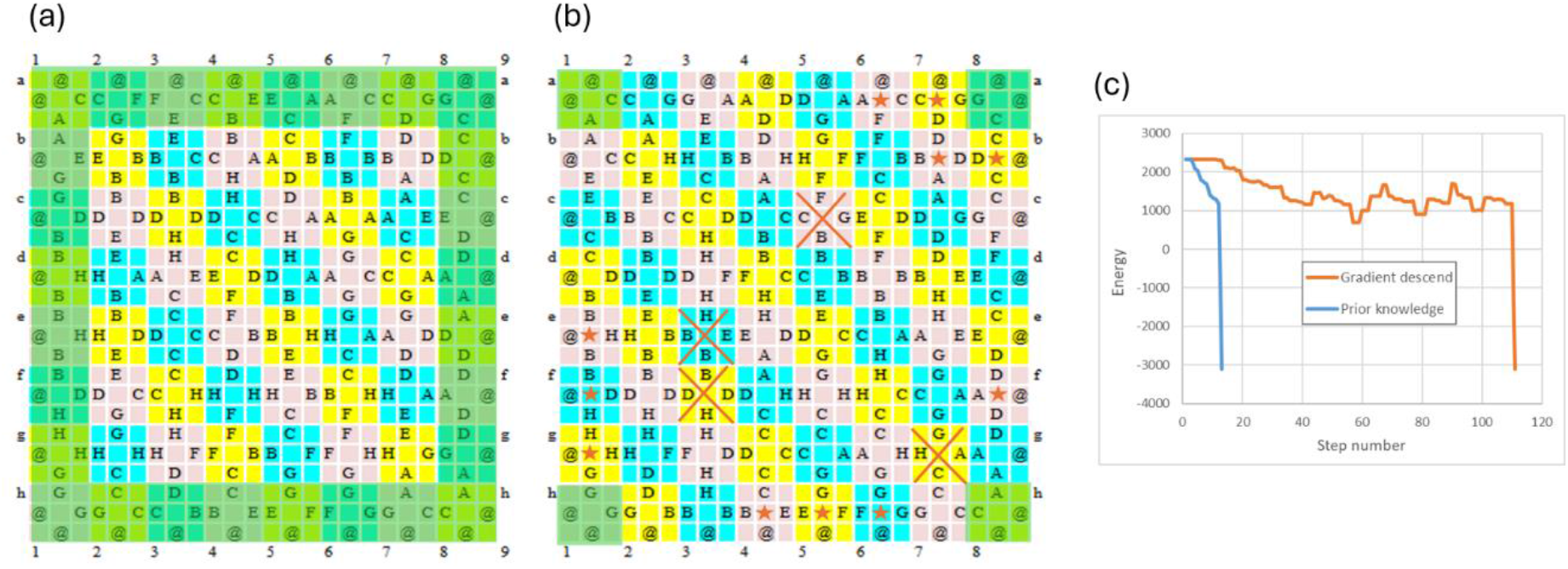
Solution of 8X8 edge-matching puzzle. (a) A correct solution obtained by the D-Wave hybrid solver based on a pre-exposed frame (marked in green). (b) A partial solution obtained based on pre-exposed corners. Orange crosses mark duplicate use of the same piece. Orange stars mark locations consistent with the full solution. (c) A solution obtained by nucleation guided by energy values (gradient descent), compared with nucleation based on prior knowledge of the solution.

The Ising energy of the complete solution is *H*=−3106.18 (after normalization of the Ising coefficients to fit the allowed range). The Ising energy of the incomplete partial solution produced by D-Wave is −3105.91, lacking only 0.005% of the total change in energy needed to reach the correct solution. This shows that a close resemblance in energy does not guarantee a close resemblance in spin configuration. Significantly complex problems set a high standard of precision for the analog components of the quantum annealer in order to obtain a unique and correct solution.

The plot in Fig. 1c shows the trace of the Ising energy calculated during a spiral nucleation process that produces a complete solution. Selection of pieces at each step was guided by steepest energy descent. However, the energy cannot decrease strictly monotonically because the process is constrained by the available matches. For comparison, selecting pieces based on prior knowledge of the solution is 8 times faster, and this factor is expected to increase with the complexity of the problem.

### B. HYBRID PIPELINE TO SOLVE 16×16 PUZZLES

For 16×16 puzzles, neither classical nucleation nor the hybrid quantum solver is sufficient on its own. Classical nucleation alone has been tested over several CPU-years, using multiple scoring strategies, yet gradient-descent-based approaches fail due to the large number of frustrated configurations (see Supplementary Material). Nevertheless, a full solution becomes attainable through a combined hybrid pipeline.

The corner pieces and their nearest neighbors are strategically critical, determining much of the subsequent solution structure. In the selected 16×16 puzzle, these 12 positions form roughly 4×10^14^ possible combinations, marked by L-shaped frames in Fig. 2a. Thus, the first step is to expose the corners and most of the frame pieces, starting from 5 pre-exposed clues highlighted in green in Fig.2b.

**FIGURE 2.**
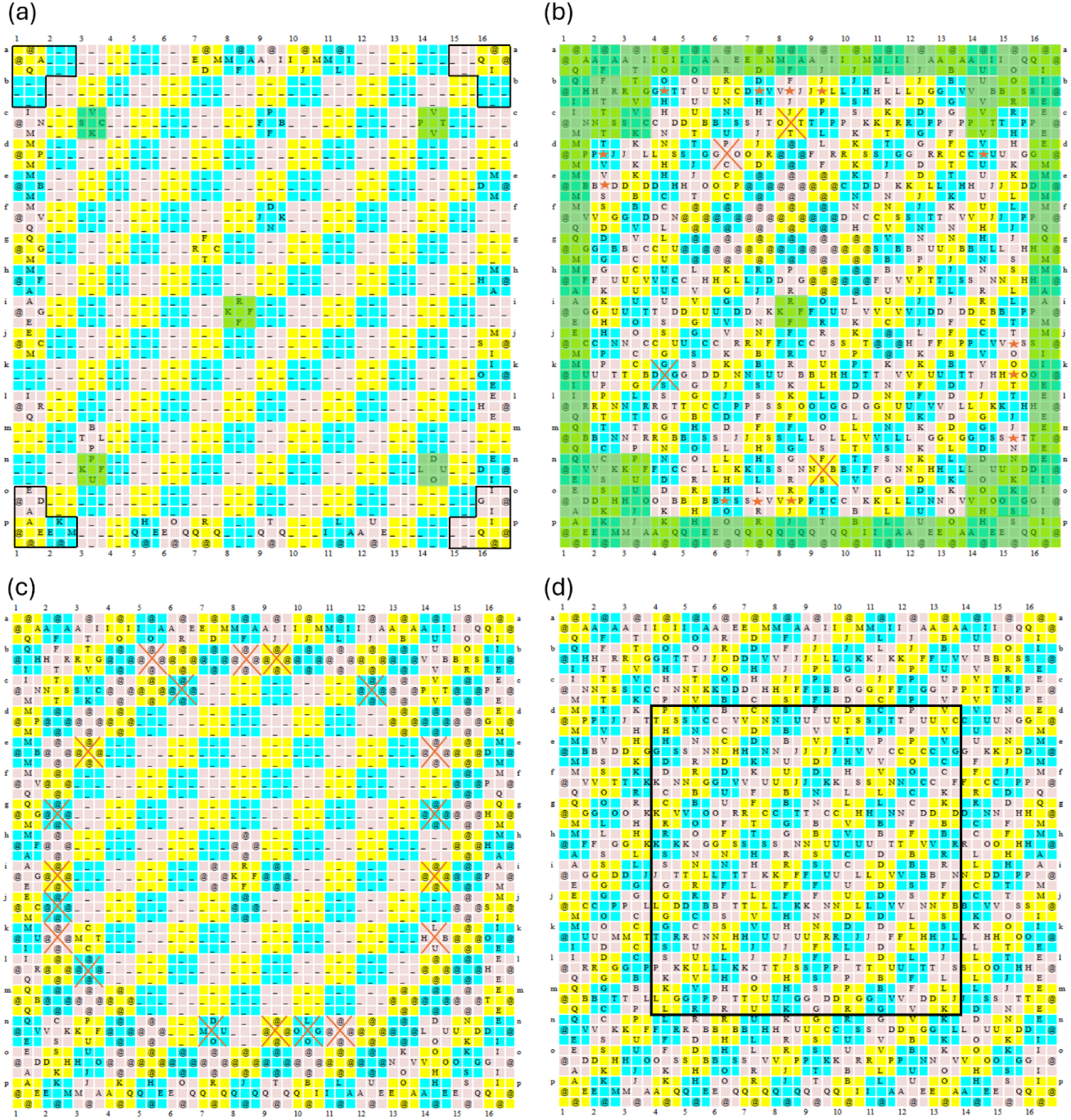
Solution steps for a 16×16 virtual puzzle with pre-exposed clues. The letter ‘@’ denotes either the frame edge or multiple options. (a) Initial step with 5 clues exposed (marked in green shade) is based on logical conjunction between partial solutions of quantum annealing with soft constraints, presenting 33 locations. (b) Quantum annealer solution with hard constraints, with pre-exposed pieces marked in green shade. Orange crosses and stars denote mismatched pieces and persistent pieces, respectively. (c) Quantum annealer solution with soft constraint, using the ancilla method. Orange crosses denote locations where a correct option is missing, determined by correlation with hard constraint solutions. (d) Complete solution obtained by nucleation from step c, after faults removed. The middle frame marks a region that is immediately recovered by deduction once lines L1–L3 are correctly determined.

Let L1 denote the first ring of tiles adjacent to the outer frame, L2 the next ring inward, and so forth. The hybrid 2p quantum annealer ran 10 minutes for the solution labeled QA7220 and 20 minutes for the solution QA7239, using soft constraints (the ancilla method as specified in Eq.8 with γ=0.050,0.015 respectively). The problem required 132,195 spins, including 251 ancilla spins. Each partial solution yielded an options matrix indicating multiple piece choices. Comparison with the known complete solution showed that QA7220 contained 71 locations with correct options (41 locations in L1), and QA7239 contained 62 locations correct (42 in L1). A major practical difficulty is determining which locations are missing the correct option, a task that can be solved only through repeated independent annealer runs. For the sake of evaluation, the logical conjunction of two option matrices was used, which was intersected with the complete solution. This produced the exposed pieces shown in Fig. 2a. The combinatorial reductions are listed in Table 1. From these exposed pieces, classical deduction can determine 90% of the pieces in L1, including the corners and their nearest neighbors.

**Table 1.** Combinatorial Reduction by Different Solvers.

| Solver | Combinatoric span in L1<br>with 5 clues pre-exposed | Combinatoric span in L2+L3<br>with L1 pre-exposed |
| --- | --- | --- |
| Setup (with deduction) | $1.3 \times 10^{98}$ | $1.2 \times 10^{123}$ |
| Quantum annealer | $2.8 \times 10^{86}$ | $6.9 \times 10^{101}$ |
| Relaxed selection waves | $3.3 \times 10^{94}$ | $1.6 \times 10^{120}$ |
| Weighted swap, 3 repeats | $9.8 \times 10^{97}$ | $5.1 \times 10^{117}$ |
| Weighted swap, AI | $5.9 \times 10^{91}$- $1.4 \times 10^{97}$ | $1.6 \times 10^{122}$ |
| Partial (60%) nucleation | NA | $7.9 \times 10^{118}$ |

For comparison, 256 relaxed partial solutions were generated using selection waves with the same five clues. On average, each solution contained 7 correct pieces in L1 and 6 in L2+L3 combined. A set of likely options was generated by selecting pieces that appeared at least 20 times across the partial solutions. To evaluate the combinatorial reduction, I cleared selections in faulty locations based on prior knowledge. The resulting combinatorial span was improved relative to the initial span, but it remained far behind the reduction achieved by the quantum-annealer solution (see Table 1).

Another effective classical approach for generating partial solutions is the weighted-swap method provided by the popular Eternity II editor [13]. 20 partial solutions with 90±2% matching edges were examined in two cases: one starting with 5 clues, and the other starting with all L1 pieces correctly placed and fixed. Compared with the complete solution, the first case showed, on average, one persistent piece in L1. Solutions starting with L1 pieces pre-exposed revealed 7 persistent pieces in L2. The AI-based analysis provided a significant improvement in the first case, although still far behind the improvement achieved by the quantum-annealer solution (see Table 1).

The AI process was found to be effective in recovering the missing 10% of pieces in L1 after the initial deduction step. Because it involves random selection of orientations, it is not deterministic and should be repeated a handful of times. One instance of the process is documented in the GitHub repository under the folder AI_example. It required generating 22 partial solutions by nucleation up to ring L3, then running the AI step with two iterations, applying logical conjunction with the initial configuration, erasing selections outside L1, and finally running nucleation of L1. The resulting pieces are the green-shaded ones shown in Fig. 2b, selected from a handful of other possible choices.

Fig.2b was generated using the D-Wave hybrid solver with hard constraint (α=0.0125 and β=0.25), starting from a pre-exposed L1 configuration (file QA7281, with 76,422 spins and 14 min run time). The energy of the solution is -56,185.8, whereas a complete solution scores -56,187.9. Most of the pieces (all but 4) are fully matched on their edges. 18 undetermined locations could not be resolved into a full nucleation. A direct benefit of the hard constrained annealing turned out to be 12 persistent pieces found in L2, in common with the complete solution.

Figure 2c shows the corresponding soft-constraint solution (file QA7050; 76,422 spins; 16.7 minutes). Most locations contain multiple candidate pieces (marked ‘@’). Faults, locations missing the correct option, are indicated by orange crosses. Most faults can be removed through comparison to the hard-constraint solution. After correcting all such locations, the combinatorial span decreases dramatically (Table 1).

Finally, starting from the corrected soft-constraint configuration, a 64-thread nucleation procedure was run. Nucleation operates on a progressively shrinking search space, as additional piece constraints propagate across the board. One thread reached the full solution after 24 hours, and another after four days. Completion occurs rapidly once enough pieces in L2 and L3 are resolved because deduction accelerates the remaining placement process.

In summary, the hybrid quantum annealer does not outperform classical solvers in isolation. However, it provides a critical advantage: it reduces the combinatorial search space sufficiently for classical nucleation to succeed. As demonstrated, even a quantum-driven reduction in combinatorial span, from 10^123^ to 10^101^, can suppress misleading search directions (“bad leads”) enough to make the full solution tractable.

A generalization of this experience suggests that a pipeline capable of achieving a complete solution to the 16×16 Eternity II puzzle should include the following steps:

1. Perform multiple hybrid quantum annealing runs using soft constraints, retain persistent selections, and apply deduction.
2. Run nucleation around the resulting configuration to generate multiple partial solutions filled up to ring L3. Recover the L1 pieces based on persistence, using AI assistance as needed. This should produce a small set of candidate L1 configurations that includes the correct one.
3. Perform additional quantum annealing runs initialized with the recovered L1 configurations, using both soft and hard constraints. Determine the L2 pieces based on persistence.
4. Run nucleation until a complete solution is obtained.

Further improvements may arise from enhancements to the hybrid quantum solvers. Techniques for mitigating precision errors have recently been proposed [14]. Additional promise lies in realization of the quantum annealer by quantum gate circuits [15], allowing for algorithmic enhancements. For example, the quantum approximate optimization algorithm (QAOA) has been shown to outperform quantum annealing in some settings [16]. Qubits could be used to encode positional addresses of pieces, reducing ∼256 possible positions to an 8-qubit representation. In principle, a quantum-gate model could expose multi-option superpositions directly. The only limitation is that commercially available systems (e.g., IBM’s Qiskit-based processors) are currently limited to roughly 100 qubits.

## IV. PROSPECT FOR SOLVING PROTEIN STRUCTURE USING HYBRID QUANTUM PIPELINE

Beyond its mathematical appeal, the analogy between the edge-matching puzzle and protein folding highlights a central question in biophysics: how do proteins reliably reach their correct three-dimensional structure, typically a minimum-energy state, despite the immense complexity of the search space? The Levinthal paradox points out that if protein folding were an NP-hard search over all configurations, nature could not achieve correct folding within milliseconds. One viewpoint suggests that residue-to-residue matching is largely unambiguous, allowing folding to proceed in a near-deterministic manner like a solving a conventional jigsaw puzzle [17], [18]. Others argue that even simplified models, such as lattice self-avoiding walks, demonstrate extreme sensitivity to initial conditions [19]. Working through the Eternity II puzzle provides a concrete analogy: the final solution depends critically on the initial configuration and the number of local frustrations encountered along the way. AlphaFold [20] is currently the most successful machine-learning tool for predicting protein structures from sequence. Nevertheless, the protein structure database contains even small peptides whose folded forms deviate from AlphaFold’s predictions (for example, entries <u>A5PJD8</u> and Q4JEI2). Foldit [21] represents another approach: it evaluates a user-proposed structure by computing its energy and applying gradient-descent refinements. However, Foldit cannot guarantee convergence to the global minimum and depends on user intuition and manual perturbations.

This suggests that protein folding indeed involves a solution to NP-hard energy minimization, a challenge as formidable as the NP-complete edge-matching puzzle. The face-matching aspect of the “end game” in protein folding [22], where the structure becomes densely packed, is perhaps one facet of the complexity. To appreciate the complexity, one can examine a far simpler problem, a self-avoiding walk on a 2D lattice without interactions. Bahi et al. demonstrated [23] that beyond 107 building blocks, certain structures cannot be altered by a single fold due to self-avoidance. Such models preserve the essential question: does solving part of a protein’s structure significantly simplify the remaining search?

Zwanzig [24], [25] provided a thermodynamic perspective on the Levinthal paradox by introducing a partition function over “correct” and “incorrect” configurations with an energetic bias favoring the native state. His model suggests that decisions between competing conformations occur in parallel—implying that the protein samples many configurations simultaneously, somewhat analogously to a quantum superposition. In this interpretation, the protein initially occupies a coherent ensemble of folded states and, through thermal relaxation and decoherence, selects conformations according to Boltzmann statistics. This explains how the system reaches lower energy states more efficiently than a classical model. Support for this idea is found in examples of open quantum systems that maintain partial coherence [26]. Hints for actual quantum process in protein folding may be found in a parallel evolution of two reaction parameters related to coiling state and hydration [27]. Another suspected quantum-like phenomenon is the allosteric effect whereby molecular interaction at one site of a protein causes a substantial conformational change at a distant site [28]. These observations raise the question: does protein folding share mechanistic similarities with quantum annealing?

Naturally occurring proteins have been shaped by evolutionary pressure to minimize frustration [29], enabling fast and reliable folding. Under such conditions, a funneled energy landscape guides nucleation toward the native structure. This has led some researchers to argue that proteins with NP-hard folding behavior would not persist in nature. Moreover, folding often involves interactions with the cellular environment or molecular chaperones; the native structure may simply be a sufficiently metastable, functional conformation. A decisive test may come from designing de-novo proteins mapped deliberately to NP-hard computational tasks and examining whether their folded structures reflect valid solutions.

Despite these uncertainties, quantum annealing has shown promise for small proof-of-concept protein-folding models. Perdomo-Ortiz and a D-Wave team [30] demonstrated structure prediction for a five-residue lattice protein by encoding directional steps through spin variables and minimizing interaction energies. The corresponding energy expression was a third-order Ising model, decomposed into several second-order models distributed across multiple quantum processing units. Subsequent advances allowed analysis of a 14-residue protein on a pure quantum annealer and up to 64 residues using a hybrid D-Wave solver [31]. Nevertheless, current assessments conclude that the technology is not yet mature for large-scale protein-folding problems [32].

Both edge-matching puzzles and protein folding can be viewed as elaborations of the traveling-salesman path problem: the path corresponds either to the polymer chain or to the sequence of correctly placed puzzle pieces. In both cases, edge values depend on the history of the path. In puzzles, an edge reflects the mismatch penalty between a candidate piece and its provisional neighbors; in proteins, it corresponds to the interaction energy between the next monomer and surrounding residues. Using a binary encoding, amino-acid coordinates can be expressed as cumulative folding decisions, each represented by a spin. These formulations lend themselves naturally to polynomial unconstrained binary optimization, which hybrid quantum-classical solvers are designed to address. It is therefore expected that future protein-structure prediction tools will integrate quantum annealing with protein-nucleation algorithms, following approaches similar to the hybrid pipeline presented in this work.

## ACKNOWLEDGMENT

D-Wave computation was financed by the faculty of chemistry in the Weizmann Institute of Science. The author is grateful for initial support by D-Wave Leap Quantum LaunchPad program. Classical computational work was carried out on the local faculty high-performance computing facility CHEMFARM, which is supported in part by the Ben May Center for Chemical Theory and Computation. The author acknowledges helpful discussions with Microsoft Copilot and with Shai Bagon and Michael Elbaum from the Weizmann Institute and with David Agard from the University of California.

## REFERENCES

[1] C. E. Leiserson, C. Stein, R. L. Rivest, and T. H. Cormen, Introduction to Algorithms, 2nd ed. Cambridge, MA, USA: MIT Press, 2001.

[2] A. Wigderson, Mathematics and Computation: A Theory Revolutionizing Technology and Science. Princeton University Press, 2019.

[3] E. D. Demaine and M. L. Demaine, “Jigsaw Puzzles, Edge Matching, and Polyomino Packing: Connections and Complexity,” Graphs and Combinatorics, vol. 23, no. S1, pp. 195–208, Jun. 2007, doi: 10.1007/s00373-007-0713-4.

[4] “Eternity two discussion forum.” [Online]. Available: https://groups.io/g/eternity2

[5] G. Harris, B. J. Vanstone, and A. Gepp, “Automatically Generating and Solving Eternity II Style Puzzles,” in Recent Trends and Future Technology in Applied Intelligence, vol. 10868, M. Mouhoub, S. Sadaoui, O. Ait Mohamed, and M. Ali, Eds., in Lecture Notes in Computer Science, vol. 10868., Cham: Springer International Publishing, 2018, pp. 626–632. doi: 10.1007/978-3-319-92058-0_60.

[6] P. Niang, “Solving the Eternity II Puzzle using Evolutionary Computing Techniques,” 2010.

[7] L. Verhaard, “EII Solver.” Accessed: Apr. 10, 2026. x[Online]. Available: https://www.shortestpath.se/eii/results.html

[8] M. W. Johnson et al., “Quantum annealing with manufactured spins,” Nature, vol. 473, no. 7346, pp. 194–198, May 2011, doi: 10.1038/nature10012.

[9] M. Booth, S. P. Reinhardt, and A. Roy, “Partitioning Optimization Problems for Hybrid Classical/Quantum Execution”, [Online].Available: https://www.dwavequantum.com/media/jhlpvult/partitioning_qubos_for_quantum_acceleration-2.pdf

[10] “OpenJij documentation.” Accessed: Jul. 30, 2025. [Online]. Available: https://tutorial.openjij.org/en/intro.html

[11] P. L. Hammer, P. Hansen, and B. Simeone, “Roof duality, complementation and persistency in quadratic 0–1 optimization,” Mathematical Programming, vol. 28, no. 2, pp. 121–155, Feb. 1984, doi: 10.1007/BF02612354.

[12] A. Power, Y. Burda, H. Edwards, I. Babuschkin, and V. Misra, “Grokking: Generalization Beyond Overfitting on Small Algorithmic Datasets,” Jan. 06, 2022, arXiv: arXiv:2201.02177. doi: 10.48550/arXiv.2201.02177.

[13] alcibiadefr, Eternity II Editor, https://sourceforge.net/projects/eternityii/. [Online]. Available: https://sourceforge.net/projects/eternityii/

[14] K. Ohno and N. Togawa, “Mitigating Precision Errors in Quantum Annealing via Coefficient Reduction of Embedded Hamiltonians,” IEEE Transactions on Quantum Engineering, pp. 1–21, 2026, doi: 10.1109/TQE.2026.3678999.

[15] D. Aharonov, W. van Dam, J. Kempe, Z. Landau, S. Lloyd, and O. Regev, “Adiabatic Quantum Computation Is Equivalent to Standard Quantum Computation,” SIAM Rev., vol. 50, no. 4, pp. 755–787, Jan. 2008, doi: 10.1137/080734479.

[16] N. Sachdeva et al., Quantum optimization using a 127-qubit gate-model IBM quantum computer can outperform quantum annealers for nontrivial binary optimization problems. 2024. doi: 10.48550/arXiv.2406.01743.

[17] S. C. Harrison and R. Durbin, “Is there a single pathway for the folding of a polypeptide chain?,” Proc. Natl. Acad. Sci. U.S.A., vol. 82, no. 12, pp. 4028–4030, Jun. 1985, doi: 10.1073/pnas.82.12.4028.

[18] A. Sali, E. Shakhnovich, and M. Karplus, “How does a protein fold? | Nature,” Nature, vol. Andrej, pp. 248–251, 1994.

[19] J. M. Bahi, N. Cote, and C. Guyeux, “Chaos of protein folding,” in The 2011 International Joint Conference on Neural Networks, San Jose, CA, USA: IEEE, Jul. 2011, pp. 1948–1954. doi: 10.1109/IJCNN.2011.6033463.

[20] J. Jumper et al., “Highly accurate protein structure prediction with AlphaFold,” Nature, vol. 596, no. 7873, pp. 583–589, Aug. 2021, doi: 10.1038/s41586-021-03819-2.

[21] F. Khatib et al., “Algorithm discovery by protein folding game players,” Proceedings of the National Academy of Sciences, vol. 108, no. 47, pp. 18949–18953, Nov. 2011, doi: 10.1073/pnas.1115898108.

[22] M. Levitt, M. Gerstein, E. Huang, S. Subbiah, and J. Tsai, “PROTEIN FOLDING: The Endgame,” Annu. Rev. Biochem., vol. 66, no. 1, pp. 549–579, Jun. 1997, doi: 10.1146/annurev.biochem.66.1.549.

[23] J. M. Bahi, C. Guyeux, K. Mazouzi, and L. Philippe, “Computational investigations of folded self-avoiding walks related to protein folding,” Computational Biology and Chemistry, vol. 47, pp. 246–256, Dec. 2013, doi: 10.1016/j.compbiolchem.2013.10.001.

[24] R. Zwanzig, “Simple model of protein folding kinetics.,” Proc Natl Acad Sci U S A, vol. 92, no. 21, pp. 9801–9804, Oct. 1995.

[25] R. Zwanzig, A. Szabo, and B. Bagchi, “Levinthal’s paradox.,” Proc. Natl. Acad. Sci. U.S.A., vol. 89, no. 1, pp. 20–22, Jan. 1992, doi: 10.1073/pnas.89.1.20.

[26] A. Buchleitner, C. Viviescas, and M. Tiersch, Eds., Entanglement and Decoherence: Foundations and Modern Trends, vol. 768. in Lecture Notes in Physics, vol. 768. Berlin, Heidelberg: Springer Berlin Heidelberg, 2009. doi: 10.1007/978-3-540-88169-8.

[27] S. Seifer and M. Elbaum, “Thermal inactivation scaling applied for SARS-CoV-2,” Biophysical Journal, vol. 120, no. 6, pp. 1054–1059, Mar. 2021, doi: 10.1016/j.bpj.2020.11.2259.

[28] H. Hofmann, “All over or overall – Do we understand allostery?,” Current Opinion in Structural Biology, vol. 83, p. 102724, Dec. 2023, doi: 10.1016/j.sbi.2023.102724.

[29] P. G. Wolynes, “Evolution, energy landscapes and the paradoxes of protein folding,” Biochimie, vol. 119, pp. 218–230, Dec. 2015, doi: 10.1016/j.biochi.2014.12.007.

[30] A. Perdomo-Ortiz, N. Dickson, M. Drew-Brook, G. Rose, and A. Aspuru-Guzik, “Finding low-energy conformations of lattice protein models by quantum annealing,” Sci Rep, vol. 2, no. 1, p. 571, Aug. 2012, doi: 10.1038/srep00571.

[31] A. Irbäck, L. Knuthson, S. Mohanty, and C. Peterson, “Folding lattice proteins with quantum annealing,” Phys. Rev. Res., vol. 4, no. 4, p. 043013, Oct. 2022, doi: 10.1103/PhysRevResearch.4.043013.

[32] T. Scheiber, M. Heller, and A. Giebel, “Exploring Quantum Annealing for Coarse-Grained Protein Folding,” Aug. 14, 2025, arXiv: arXiv:2508.10660. doi: 10.48550/arXiv.2508.10660.

