## Supplementary material for "A Pipeline for Solving Edge-Matching Puzzles and Their Implications for Protein Folding"

Shahar Seifer

### 1. Ising model for matching two groups of pieces with a matching edge

We allow for the same number of down spins in each group, aiming for the cases of either zero selection in each group or one selected in each group.

$$H = \lambda\left[\frac{|A| - \sum_{i\in A} S_i}{2} - \frac{|B| - \sum_{i\in B} S_i}{2}\right]^2 =$$

$$\frac{\lambda}{4}[|A| - |\mathrm{B}|]^2 + \frac{\lambda}{4}\left[-2|A|\sum_{i\in A} S_i \; - 2|B|\sum_{i\in B} S_i + \left(\sum_{i\in A} S_i\right)^2 + \left(\sum_{i\in B} S_i\right)^2\right]$$

$$-\frac{\lambda}{2}\left[-|A|\sum_{i\in B} S_i \; - |B|\sum_{i\in A} S_i + \left(\sum_{i\in A} S_i\right)\left(\sum_{i\in B} S_i\right)\right] =$$

$$\frac{\lambda}{4}[|A| - |\mathrm{B}|]^2 + \left(\sum_{i\in A} S_i - \sum_{i\in B} S_i\right)\frac{\lambda}{2}(|B| - |A|)$$

$$+\frac{\lambda}{2}\sum_{i<j\in A} S_i S_j + \frac{\lambda}{2}\sum_{i<j\in B} S_i S_j - \frac{\lambda}{2}\sum_{i\in A, j\in B} S_i S_j$$

### 2. Ancilla method for softening a constraint in the Ising model

Suppose we wish to add a constraint for one spin down ($S_i = -1$) out of M spins. A standard way is to add the term

$$\Delta H = \left[\sum\nolimits_{i=1}^{M} S_i - (M-2)\right]^2$$

However, there is a sever penalty due to the quadratic contribution (up to 36 if *M*=4). A quadratic contribution is the only way to add a positive penalty in the Ising model.

A softer penalty can be achieved by splitting the M spins into two equal groups and adding an ancilla spin $S_{anc}$ that helps settle the penalty to minimum when there is only one spin down out of the two groups. The following term achieves this constraint

$$\Delta H = \tfrac{M+\delta}{4}\left(\sum\nolimits_{j=1}^{\lceil \frac{M}{2} \rceil} S_j - \sum\nolimits_{j=\lceil \frac{M}{2} \rceil+1}^{M+\delta} S_j - 2S_{anc}\right)^2 - \sum\nolimits_{j=1}^{M+\delta} S_j$$

$$\delta = \begin{cases} 0, if\ M\ is\ even \\ 1, if\ M\ is\ odd \end{cases}$$

For example, in the case *M*=4 the maximum penalty is 8 and is -2 at the correct minimum.

### 3. Ising model formalism in the traveling salesman path problem

The salesman's options are represented by a spin $S_{t,j}$ that is -1 if the salesman visits node *j* at time $t$, and is +1 otherwise. The Ising model is written

$$H_{TSPP} = \sum_{t=1}^{N}\sum_{i=1}^{N}\sum_{j=1}^{N}\frac{1}{4}d_{ij}(1 - S_{i,t})(1 - S_{j,(t+1)modN}) \qquad \text{(S1)}$$

The solution involves only one spin down at a particular location or time, so the following two constrained optimizations, ideally equal to zero, are added to the score function $H$ with Lagrange multipliers:

$$\Delta H = \lambda_1(\textstyle\sum_{j=1}^{N} S_{t,j} - N + 2)^2 + \lambda_2(\textstyle\sum_{t=1}^{N} S_{t,j} - N + 2)^2 \qquad \text{(S2)}$$

The task may be implemented in a classical solver such as OpenJij; however, it does not necessarily provide an exact solution within a reasonable time. A specifically tailored "selection wave" approximation method is described in the main text and demonstrated in a python code.

### 4. Failure of classical gradient descent in 16x16 puzzle

How useful is it to guide a nucleation solution approach by a score function?

The options matrix $\bar{\bar{O}}$ (of size 256x1024) determines for each position out of 256 and each available piece and orientation if the option is still valid (1) or eliminated (0). We consider a "virtual" 16x16 puzzle that resemble the Eternity II puzzle, and accordingly the starting configuration is set with 5 predetermined pieces (the center piece at location i8 and 4 clues at c3, c14, n3, and n14). We also predetermine the frame pieces to be present only in the frame positions. We allow also the correct corner pieces to be present at start. Then a deduction operation is activated, which eliminates further options as described in the Methods section of the main text.

Several pseudo score functions are calculated beside the Ising model energy. One is the rank of the matrix $\bar{\bar{O}}$, which should reach 256 in the final solution where all the positions have exactly one candidate piece.

$$Score_1(\bar{\bar{O}}) = Rank(\bar{\bar{O}})$$

The "algebraic score" is the sum of elements in a vector $v$ that is the least square solution of the following set of linear equations (that should also reach 256 in the final solution)

$$\bar{\bar{O}}\,v = \begin{bmatrix} 1 \\ \vdots \\ 1 \end{bmatrix}, \quad Score_2(\bar{\bar{O}}) = \textstyle\sum_i v_i$$

The “optimistic score” counts how many positions still hold a valid choice from the complete solution that is known in advance. The plots in Fig.S1 are traces of attempts to complete a spiral nucleation from the frame.

Fig.S1a is a case in which the solution is guided by the algebraic score function. It succeeds up to step 11 to approach correct values in rank and algebraic score. However, further steps decrease the optimistic score to zero, and carries with it the other scores.

Fig.S1b demonstrated nucleation guided by a full Ising model energy calculation. The progress halts after initial partial success. Searching the global energy minimum based on local energy gradients resembles walking in a maze of high dimensionality. The path is obstructed by too many confinements, making very low expectancy for timely success.

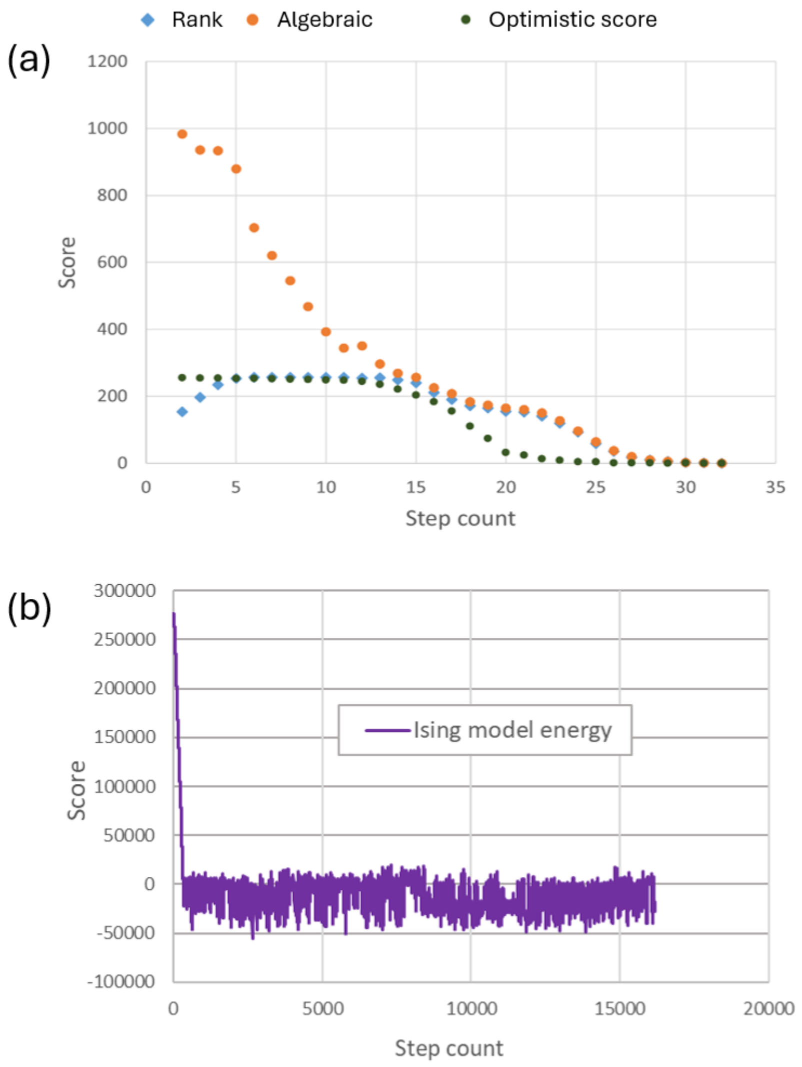


*Figure S1. The progress of 4 score functions in a gradient descent attempt to solve a virtual 16x16 puzzle. (a) Guided by algebraic score the progress halts at step 13. (b) Guided by the Ising model energy the solution does not progress beyond initial partial success.*
